# Stochastic motility-adhesion switch within the *E. coli* K-12 strains revealed by gene expression clustering

**DOI:** 10.64898/2025.12.14.694247

**Authors:** Anna D. Kaznadzey, Tatiana A. Bessonova, Uliana D. Kuznetsova, Mikhail S. Gelfand, Maria N. Tutukina

## Abstract

Biofilm formation in *Escherichia coli* arises from regulatory programs that can diverge even among genetically identical cultures. Exploiting spontaneous replicate-to-replicate variation in *E. coli* K-12 MG1655, we mapped expression modules whose behavior tracks biofilm abundance across four genetic backgrounds: wild type; strain with deleted gene for hexuronate regulator UxuR Δ*uxuR*; an *uxuR* translation-disrupted retaining locus-derived sRNAs (Δ*uxuR*-tr), and strain with deleted gene for global carbon regulator cAMP-CRP (Δ*crp*). In the wild type, regulator EcpR/MatA demonstrated the highest correlation with the biofilm intensity, and its role was, in particular, seemingly reinforced by stress/pH-homeostasis and c-di-GMP-linked factors. Δ*uxuR* exhibited the strongest stochasticity in biofilm formation, while in Δ*uxuR*-tr, replicate dispersion was mostly muted, indicating that *uxuR*-derived sRNAs buffer motility-adhesion switching. In Δ*crp*, *pgaA* was de-repressed due to the absence of CsrA, with expression being highly consistent with intensity of biofilm formation, alongside stress/ion-homeostasis genes. This is in line with the suggestion that CRP regulates biofilms and motility indirectly via regulation of more local transcription factors. Overall, replicate-resolved clustering converts apparent “noise” into structured regulatory patterns, which include genes whose role in biofilm formation was not yet evident, and helps identify switch nodes between planktonic and biofilm bacterial states.

## Introduction

Bacterial biofilms are multicellular communities embedded in an extracellular polymeric substance matrix, principally polysaccharides, proteins, extracellular DNA and lipids, that confers emergent properties distinct from planktonic growth, including altered gene-expression programs and stress tolerance [1]. Within mature biofilms, microscale chemical gradients and lineage/stochastic effects generate pronounced physiological heterogeneity, with subpopulations differing in metabolism, stress responses, and growth state [2]. Biofilms are clinically and industrially consequential: in medicine, they are a common basis of device-associated and chronic infections, with *Escherichia coli* prominently implicated in catheter-associated urinary tract infections and other persistent conditions involving, for example, enteropathogenic strains [3–5]. Biofilm growth also contributes to antimicrobial tolerance and recalcitrance [6].In industry, unwanted biofilm growth (“biofouling”) burdens water treatment and distribution systems, membranes, pipelines, and marine infrastructure, incurring substantial operational and maintenance costs [7].

When bacteria switch their lifestyle from a “solitary” planktonic existence to a biofilm, a life in a community, significant changes occur in cellular morphology, physiology, and metabolism [8–11]. Most of the systems responsible for flagella formation are switched off, and the genes for proteins necessary for adhesion are activated. The transcription of *Escherichia coli* genes whose products are involved in flagella formation is controlled by the motility sigma factor, σ28, FliA, and the master regulator FlhCD [12,13]. However, some cell motility must still be maintained: deletion of the gene encoding FliA led to changes in the architecture of biofilms formed by E. coli K-12 MG1655 and a decrease in their thickness [14].

The common *mat/ecp* fimbriae contribute to adhesion and early biofilm formation on abiotic and host surfaces [15]. Their transcription is regulated by the LuxR-family factor MatA (also called EcpR), which activates the *mat* operon and simultaneously represses the flagellar master regulator *flhDC*, thereby switching cells from a motile to a sessile state [16]. MatA is embedded in higher-order regulatory circuits involving RcsB and H-NS that coordinate envelope stress responses and motility repression, further reinforcing the transition toward surface attachment [17]. In parallel, small regulatory RNAs (sRNAs) such as MicA, MicF, and others modulate outer membrane composition, flagellar components, and adhesin expression, adding a post-transcriptional layer to the control of motility and biofilm formation [18,19].

Curli pili play a key role in the initial stages of adhesion to solid surfaces and subsequent intercellular interaction. The genes encoding the respective proteins are organized into two divergent operons – *csgBA* encodes the structural components, and *csgDEFG* encodes the pili assembly proteins and transport proteins [20]. The expression of both operons is under the control of the local regulator CsgD [21] from the FixJ/LuxR family and modulates the expression of not only the *csg* operon genes but also other genes encoding proteins necessary for the cell’s adaptation to life in a biofilm [22–24]. For example, it activates the transcription of *adrA* (*dgcC*), which encodes diguanylate cyclase for the synthesis of cyclic di-GMP, a secondary messenger that plays an important role in regulating many processes in biofilm life [25,26]. Activation of *adrA*, in turn, leads to increased cellulose production [27]. In addition, CsgD regulates the mRNA level of the *yoaD* gene, which encodes one of the cyclic di-GMP phosphodiesterases [23], suggesting that CsgD is involved in controlling the intracellular level of cyclic di-GMP [28,29]. By activating the transcription of *glyA* (encoding hydroxymethyltransferase), CsgD accelerates the synthesis of curli pili, which contain more glycine than other *E. coli* proteins [22].

An important component of the biofilm is the polysaccharide matrix, which includes, among other things, colanic acid and cellulose [8,27,30], which must be synthesized simultaneously with the pili. The *pgaABCD* operon is necessary for the synthesis of an adhesin – a cell-binding polysaccharide rich in hexosamines (a linear polymer consisting of β-1,6-N-acetylglucosamine residues), which is repressed by the carbohydrate metabolism regulator CsrA, which, in turn, is repressed by small regulatory RNAs CsrB and CsrC [31]. Accordingly, overproduction of CsrC leads to enhanced biofilm formation and reduces motility [32].

The synthesis of curli fimbriae and cellulose is strictly controlled at several levels by the transcription factors CsgD, CsrA, and MlrA, cyclic di-GMP, and ncRNAs. In addition to CsrB-CsrC, a number of other ncRNAs are involved in the control system, namely ArcZ, DsrA, RprA, McaS, OmrA/OmrB, GcvB, and RydC. At the top of this cascade is the stress sigma subunit RpoS, the expression of whose gene is also controlled by ncRNAs, namely ArcZ, DsrA, and RprA, primarily by altering the mRNA structure and protecting it from degradation by RNases [19].

6S RNA (SsrS) directly binds to another sigma factor, σ70, RpoD, and the β/β′-subunits of RNA polymerase. It accumulates when the cell transitions to the stationary growth phase and is able to repress the expression of genes transcribed from σ70-dependent promoters [3]. Using RNA sequencing we previously showed that SsrS is overproduced in cells within a biofilm, and its biofilm activatory function was further confirmed in physiological tests (Bessonova et al., Molecular Biology, N4, 2026 in press).

Environmental stress responses, particularly acid and oxidative stress resistance, are also tightly intertwined with *E. coli* biofilm physiology. Yet the extent to which these stress response modules shape biofilm heterogeneity among genetically identical populations remains to be clarified.

A less explored regulatory axis involves hexuronate metabolism and the GntR-family transcription factors UxuR and ExuR, hence ExuR and D-glacturonate was shown to be critical for mouse intestine colomization [33]. These paralogous regulators coordinate the utilization of hexuronic acids such as D-glucuronate and D-galacturonate, influencing central carbon and cell wall precursor fluxes [34]. Recent work further demonstrated that the *uxuR* locus gives rise to multiple sense and antisense RNAs from divergent promoters in its 3’ region, suggesting a complex regulatory architecture where UxuR protein and associated small RNAs may exert distinct functional roles [35]. Hexuronate availability has been reported to modulate biofilm formation in *E. coli* K-12 MG1655, pointing to a functional connection between this metabolic pathway and adhesive behavior [36]. However, the contribution of UxuR itself and of its non-coding RNA products to biofilm regulation in MG1655 has not been systematically dissected.

Over the last few years, it has become increasingly clear that biofilm communities are not homogeneous but instead exhibit extensive transcriptional and phenotypic heterogeneity at the level of individual cells and microcolonies [37]. Single-cell and spatially resolved studies in *E. coli* have revealed distinct physiological subpopulations that differ in metabolic activity, stress responses, and regulatory states [38]. For example, Besharova et al. demonstrated that submerged static biofilms of *E. coli* W3110 contain spatially segregated curli-producing, motile, and σ^S-active subpopulations, reflecting structured diversification during biofilm development [37]. Intra-strain sectoring has also been observed in uropathogenic *E. coli* (UPEC) colony biofilms, where “peppermint” sectors differ in rugosity and matrix composition and were linked by sequencing to emergent mutations (e.g., *nlpI*) [39].

Foundational single-cell work studied expression variability, establishing noise as an inherent driver of phenotypic spread, and explaining how feedbacks and thresholds enable bistable switching [40]. Second-messenger networks further amplify diversification: c-di-GMP signaling, implemented by numerous diguanylate cyclases and phosphodiesterases, operates through combined local and global modes, generating cell-to-cell variability that toggles motility and adhesion modules during early biofilm assembly [41]. Engineered biofilm reactors likewise show that small early differences can push identically operated systems onto divergent trajectories; microbial succession alternates between deterministic and stochastic phases, and biofilm thickness shifts the balance toward greater β-diversity across replicate carriers [42,43].

In *E. coli*, most prior work interrogates within-biofilm heterogeneity across mutants/conditions rather than replicate-to-replicate divergence in an otherwise constant background [37]. Conceptually, such divergence could reflect stochastic or history-dependent activation of decision nodes that gate the motile-adhesive switch: global carbon status via cAMP–CRP (affecting curli/cellulose through *csgD* and modulating CsrB/C), the MatA/EcpR axis that promotes *ecp/mat* fimbriae while repressing *flhDC*, hexuronate regulators UxuR/ExuR (and uxuR-derived sense/antisense RNAs) influencing motility and envelope-linked functions, acid-resistance hierarchies that intersect with biofilm circuits, c-di-GMP supermodules that partition cells into motile vs adhesive states and other mechanisms. Systematic mapping of transcriptome modules correlated with spontaneous replicate-level differences in biofilm abundance has not been performed in a K-12 MG1655 background.

Here, we address this gap by combining replicate-level phenotyping with RNA-seq and expression-pattern clustering across MG1655 and regulatory mutants (Δ*crp*, Δ*uxuR*, and a translationally disrupted *uxuR* gene). By sequencing biofilm-rich and biofilm-poor replicates grown under identical conditions, we identify transcriptional clusters whose replicate-specific behavior tracks biofilm abundance, thereby revealing regulatory, metabolic, and stress-linked pathways that bias clonal *E. coli* populations toward biofilm-prone vs biofilm-averse states.

## Materials and Methods

### Strains and growth conditions

Here, *E. coli* K-12 MG1655 [44] was used as a parent strain. The *uxuR* gene was disrupted in *E. coli* BW25113 using recombineering technique [45]. The gene::kan mutations were transferred by P1-transduction into *E. coli* K-12 MG1655. *E. coli* K-12 MG1655 *Δcrp* was obtained the same way from M182 strain [46] and *E. coli* K-12 MG1655 *ΔuxuR_tr* was constructed using gene doctoring [47].

Purified colonies were grown aerobically in Luria-Bertani (LB) medium for 16 hours at 37℃ under constant shaking. Three overnight cultures from three independent colonies were made for each strain, and each culture was then tested for biofilm formation in triplicate, so at least nine replicates were made for each strain. The overnight cultures were then diluted 1:20, and the biofilms formation was allowed for 72 hours in 96-well plates (83.3924.005, Sarstedt, Germany) at 37℃ and microaerobic conditions as described earlier [36]

### Biofilm staining and RNA extraction

From each well, part of the biofilm cells was taken for RNA extraction, part was stained with crystal violet (Sigma, USA) as described earlier [48]. Biofilm formation intensity was measured at λ=570 nm on the BioTek Synergy H1 plate reader (USA).

RNA was extracted with the TriZol reagent (Invitrogen, USA, 800 ul per well) using standard protocol and then treated with the DNAse I (Promega, USA) for 1 hour at 37℃ followed by inactivation. Resulting RNA concentration was measured on the Qubit 4 fluorometer (Thermo, USA) using the RNA HS kit (Thermo, USA).

### Quantitative PCR and sequencing

50 ng of RNA was taken in the reaction of reverse transcription made with the M-MulV RevertAid reverse transcriptase (Thermo, Lithuania) and gene-specific primers according to the manufacturer’s protocol. Then, the cDNA was used for quantitative PCR (qRT-PCR) performed with the qPCR-HS SYBR mix (Evrogen, Russia) on a DT-Prime thermocycler (DNA-Technology, Russia). Primers used for reverse transcription and amplification are listed in Table S1. No PCR products were detected in negative controls in the absence of reverse transcriptase. Data obtained from at least six statistical replicates were calculated by the dCt method and presented as the heatmap created in R.

**Table S1.**
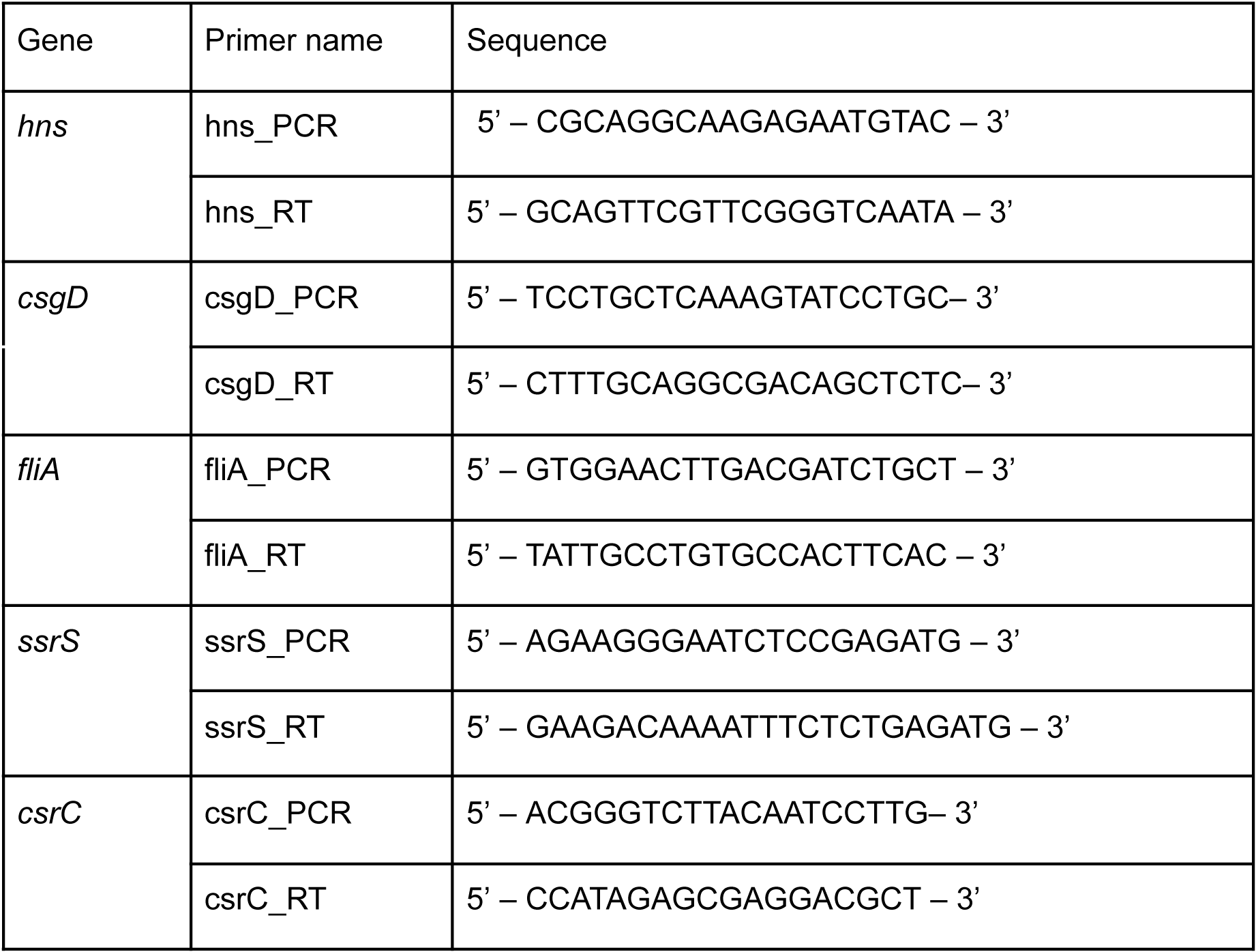
Primers used for qRT-PCR.

Based on the biofilm formation intensity and the qRT-PCR data, 4-6 replicates of each strain were chosen for whole transcriptome sequencing.

Libraries were prepared using the NebNext Ultra II directional RNA library kit (New England Biolabs, USA), quantified on the Qubit-4 and Bioanalyzer, and sequenced on the Illumina NextSeq500 as 75+75 PE in the Skoltech Core Genomics and Biomaging Facility.

### Data analysis

RNA-seq data from biological replicates of *Escherichia coli* strains were analyzed to identify clusters of genes with shared expression patterns. Raw gene count matrices were optionally normalized to Transcripts Per Million (TPM) to correct for both sequencing depth and gene length differences. Log-transformed counts (log₁(x + 1)) were then standardized across replicates using z-score normalization on a per-gene basis, ensuring comparability of expression trends across genes regardless of their absolute abundance.

For unsupervised clustering, hierarchical clustering was performed using Ward’s minimum variance method [49] and Euclidean distance as the dissimilarity metric. Gene clusters were defined by cutting the resulting dendrogram at a fixed distance threshold (d = 8.0), and within each cluster, average z-scores were computed to extract characteristic replicate-level expression patterns.

To assign gene-level pattern annotations, log₂ fold change was computed relative to the median expression per gene. Genes showing strong deviations (log₂FC > 0.7 or < –0.7 for TPM-normalized data) were marked as “high” or “low” in a given replicate.

Gene functions were annotated by mapping gene names to curated functional descriptions from EcoCyc [50], UniProt [51], and literature-based annotations.

Data analysis was performed using Python with the pandas library for data manipulation [52], NumPy for numerical computation [53], SciPy for hierarchical clustering [54], matplotlib for static graphics [55], and seaborn for clustered heatmaps [56].

## Results and discussion

### Replicate-Level Variation Reveals Spontaneous Biofilm Heterogeneity Across Strains

Four genotypes were assayed in 96-well format: wild type (wt), Δ*crp*, Δ*uxuR*, and a translation-disrupted *uxuR* allele that preserves potential sRNA transcription (Δ*uxuR-*tr). We first quantified expression levels of four biofilm-relevant regulators by qPCR, FliA, (flagellar class-3 sigma factor), CsgD (curli/matrix regulator), and two regulatory RNAs, CsrC sRNA (antagonist of CsrA), and SsrS/6S RNA (σ^70 holoenzyme modulator that was also shown to activate biofilms), and selected over- and under-expressing wells, in parallel measuring the biofilm formation intensity in each well. The results are shown in Fig. 1. We tried to correlate the intensity of biofilms with the expression levels of genes coding for the motility (*fliA*) and biofilm formation (*csgD*) regulators (Fig.1) within colonies. However, we do not observe a clear correlation, except for very low biofilm efficiency in the replica with high *fliA*-mRNA levels (A5 in *ΔuxuR*). Moderate levels of *fliA* expression is characteristic for almost all colonies, even with high biofilm intensity, since it was shown that cellular motility apparatus is apparently required for normal biofilm formation [14]. For further RNA sequencing we took colonies with the most diverse biofilm levels, including those with similar expression patterns. The same strategy was applied for the wild type cells.

**Fig. 1.**
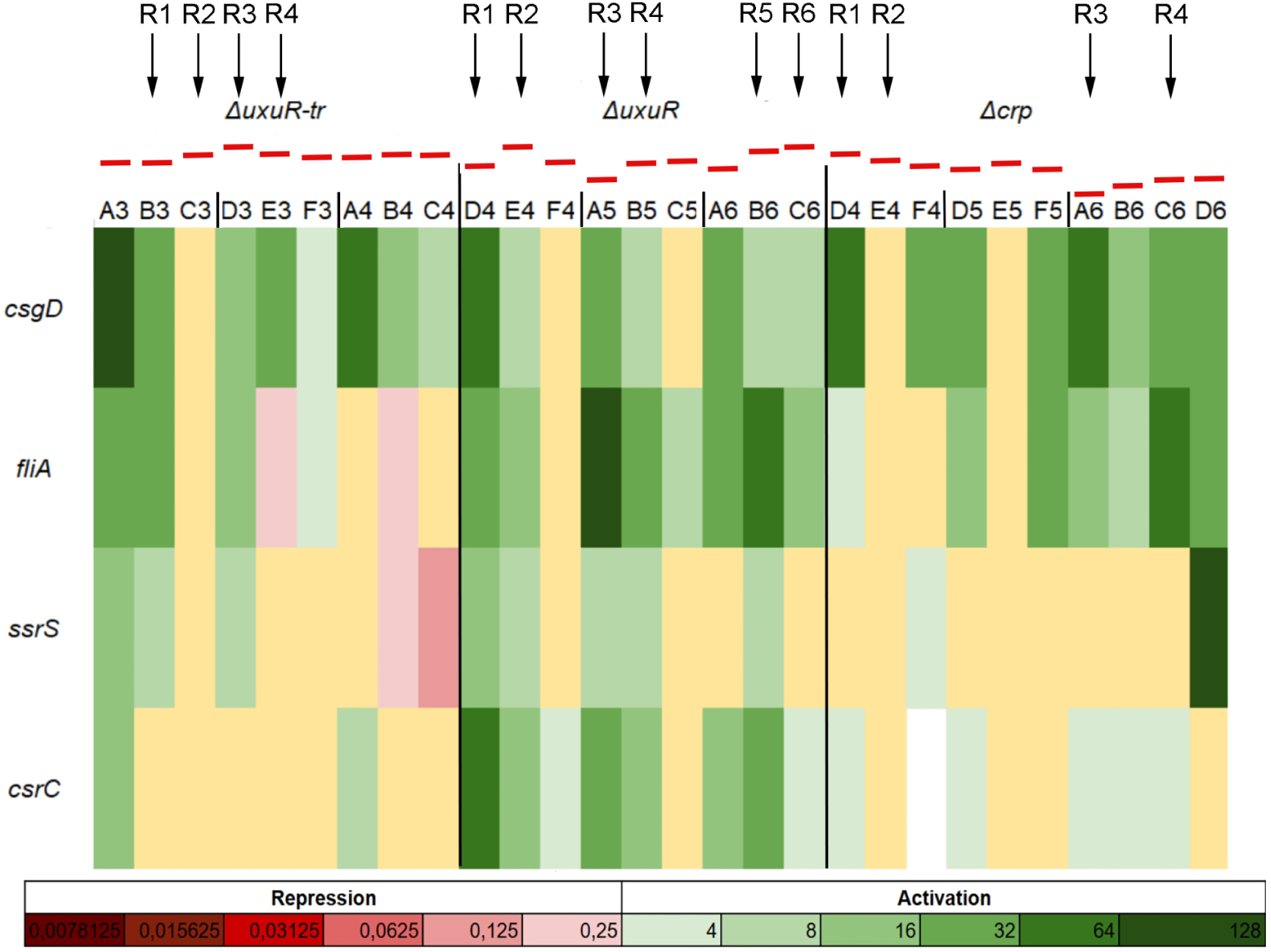
Heatmap reflecting the qPCR results as relative expression to the wild type (averaged for all replicates). Strains and well numbers are indicated above, colonies are separated by black lines. The levels of biofilm formation is indicated above the heatmap as red rectangles.

Gene expression was then grouped into replicate-specific patterns, and clusters whose signatures positively or negatively tracked biofilm abundance were prioritized for interpretation. Across strains, wt and Δ*crp* exhibited pronounced replicate-level variation in biofilm formation, with Δ*uxuR* showing even greater dispersion. In the sections that follow, we analyze each strain separately, highlighting cluster modules whose replicate behavior correlates with biofilm levels and nominating both established and under-recognized genes potentially involved in biofilm formation.

### Wild-Type Biofilm Heterogeneity Is Driven by Dual Control of Motility and Adhesion Pathways

In the wt strain 6 replicas were analyzed, yielding 14 clusters of genes with similar expression patterns among replicas (R). Biofilm abundance was highest in R1 and R2 and lowest in R3 and R4 (Fig. 2). Among the expression modules, Clusters 1, 2, 6, and 13 showed the strongest correlation with biofilm intensity. Cluster 6 was most consistent with an adhesion/matrix-on, motility-off program. Notably, it contains *ecpR*/*matA*, the LuxR-family regulator that both activates the *ecp/mat* fimbrial operon and represses the flagellar master regulator *flhDC*, thereby biasing cells toward a sessile state; MatA also interacts (directly or via higher-order circuits) with additional motility loci [57]. This nominates the MatA axis as a principal driver of the wt replicate divergence. Interestingly, this gene has not been previously reported as the one to be involved in stochastic processes related to biofilm formation.

**Fig. 2.**
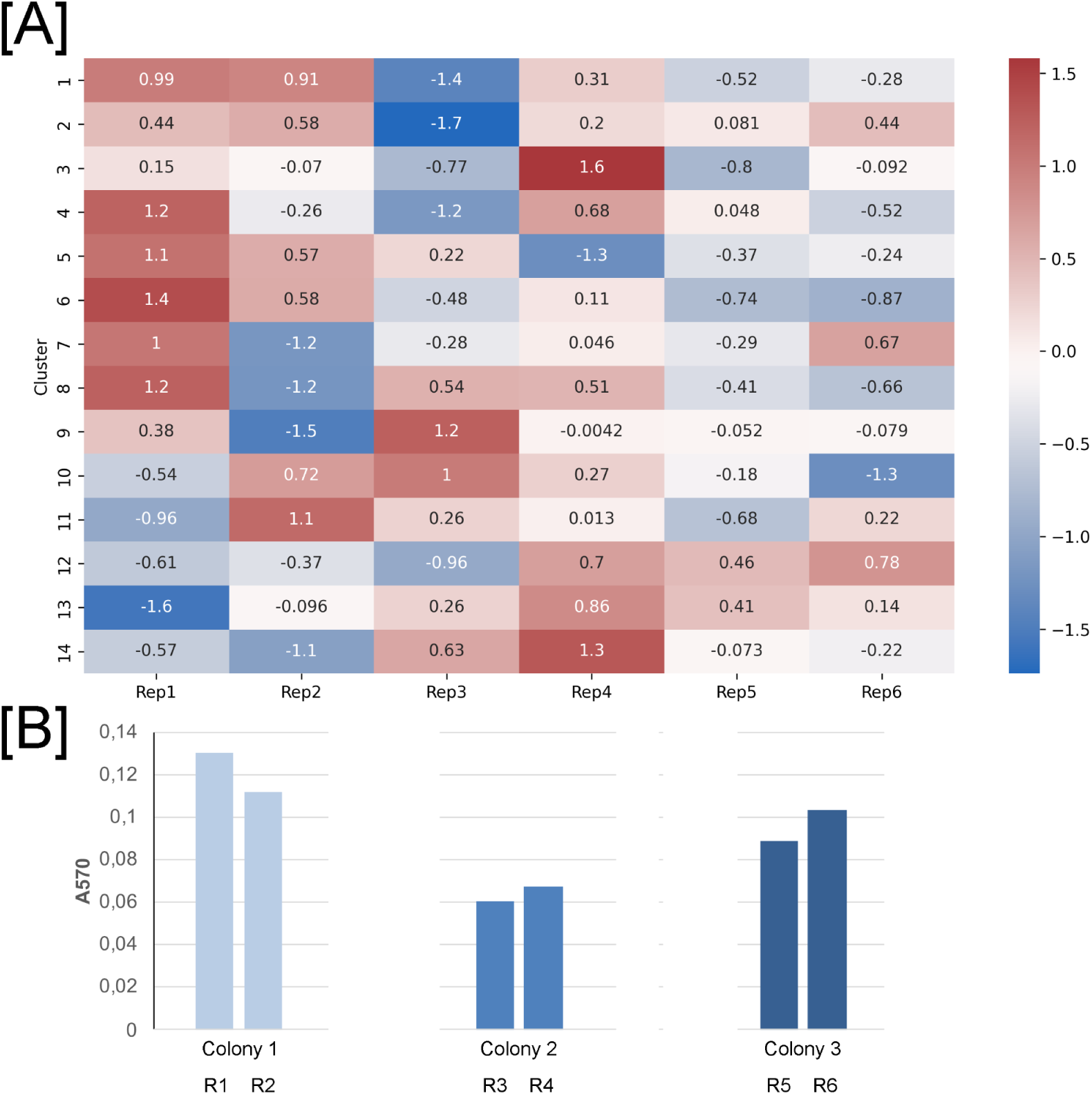
[A] Heatmap of gene expression pattern clusters for wild type *E. coli* K-12 MG1655. Values indicate mean Z-scores, the respective color scale is shown on the right. [B] Biofilm intensity formation for respective replicas.

Cluster 6 also harbors genes that plausibly reinforce matrix production and stress-tolerant states typical of biofilm-rich replicas. *nfrB* encodes a c-di-GMP–regulated glycosyltransferase that synthesizes a secreted exopolysaccharide (EPS) required as the primary receptor for phage N4; this EPS hampers motility and contributes to surface encasement, directly linking c-di-GMP signaling to matrix output[58]. *ydeO*, an AraC-family regulator upstream of *gadE* in the AR2 acid-resistance pathway, connects acid stress to adhesin regulation and has been implicated in biofilm behavior under acidic conditions, providing a route by which microenvironmental pH can tune biofilm propensity [59]. The toxin–antitoxin kinase HipA is a canonical persistence factor; perturbations of *hipA* increase multidrug-tolerant persisters and have been tied to biofilm tolerance, consistent with enrichment of slow-growing, stress-tolerant subpopulations in biofilm-high replicas [60].

Stress- and envelope-linked factors in this cluster further align with the biofilm phenotype. *yhcN/bhsA* (RpoS-associated cell-envelope stress protein) acts as a negative regulator of excessive surface adhesion; *yhcN* deletion increases biofilm and aggregation, while *yhcN* is often upregulated in biofilms, an apparent feedback that tempers over-adhesion yet is frequently observed in biofilm transcriptomes [61]. *nhaA* (Na⁺/H⁺ antiporter) together with the LysR regulator NhaR links ionic/pH homeostasis to PNAG synthesis: NhaR directly activates the *pgaABCD* operon required for PNAG production, and *nhaA/nhaR* circuits have been integrated into biofilm regulatory models, supporting a role for salt/alkali conditions in modulating matrix output within replicas [62].

Several additional genes merit cautious consideration. *murR* (MurNAc catabolism regulator) could, in principle, intersect with envelope remodeling that influences biofilm architecture, but we found no direct evidence linking *murR* to *E. coli* biofilm traits. *higB* (HigBA TA toxin) has not been singled out in MG1655 biofilms, though clinical-isolate surveys associate TA loci broadly with strong biofilm phenotypes in *E. coli* (gene-set level signal rather than *higB*-specific) [63]. *yeaH* (of the *yeaG–yeaH–yeaI* stress module) participates in nitrogen-starvation adaptation and is connected to stringent-response/persister formation, suggesting a route by which micro-nutrient differences across replicas could reinforce biofilm-prone states [64]. Finally, *trpE* sits upstream of indole production (via *tnaA*); indole is a well-established anti-biofilm signal in *E. coli*, so flux through tryptophan synthesis can indirectly modulate biofilm propensity [65].

Other gene clusters with over-expressed genes in high biofilm replicas are Cluster 1 and Cluster 2. The first one contained genes consistent with biofilm maturation and regulatory cross-talk between adhesion, stress, and efflux. Notably, *sslE*, a T2SS-secreted M60 metalloprotease—has been shown to promote biofilm maturation, polymerizing under acidic microenvironments within mature biofilms and binding eDNA/cellulose to cross-link the matrix, providing a mechanistic route to late-stage stabilization [66]. The small protein AriR (YmgB) functions as a biofilm regulator whose deletion/overexpression perturbs biofilm formation and motility and interfaces with indole signaling and acid-stress responses, indicating capacity to shift the motile↔adhesive balance under environmental cues [67]. *ecpC* encodes the usher of the *E. coli* common pilus (ECP); ECP contributes to adherence and biofilm formation across diverse *E. coli* backgrounds, supporting an early-adhesion role for this module. This complements the previous finding concerning *ecpR*/*matA* from Cluster 6 [68]. The presence of *marC* links Cluster 1 to the mar regulon context: while direct biofilm functions of *marC* remain under-defined, the regulon’s transcription factor MarA is known to regulate the *ycgZ*–*ymgABC* region (AriR locus) and thereby modulate biofilm formation, suggesting regulatory coupling between multidrug-resistance circuits and surface lifestyles [69]. Finally, *mdtA* (a component of a multidrug efflux pump) aligns with reports that specific MDT deletions alter *E. coli* biofilm growth and fitness, especially under antimicrobial stress—consistent with efflux tuning of matrix/adhesion phenotypes [70].

Cluster 2 was enriched with genes for motility/early-attachment and type II secretion functions, led by *flhA* (flagellar export apparatus) and *fliD* (flagellar cap). Classic genetics show that disrupting flagellar assembly or motility impairs *E. coli* K-12 biofilm initiation, placing *flhA/fliD* squarely in the “required-for-early-attachment” class [71]. The cluster also includes *gspM*, a structural component of the type II secretion system (T2SS); in enteropathogenic *E. coli* (EPEC), an intact T2SS and its lipoprotein substrate SslE are required for biofilm formation and virulence, and T2SSs more broadly deliver matrix-active proteins that shape biofilms [72].

Cluster 14, on the other hand, comprised genes of central biosynthesis and energy generation that are consistently downregulated in high-biofilm replicates and upregulated in low-biofilm replicates, indicating a growth-associated program that is curtailed during biofilm formation. Specifically, nucleotide and translation/replication functions (*carA*, *carB*, *codA*, *codB*, *rplC*, *rplW*, *glnW*, *glnU*, *priB*) follow the expected reduction of protein synthesis and DNA replication in biofilm mode, a hallmark of mature *E. coli* biofilms and mixed-species biofilms under microenvironmental constraints [11]. Aerobic respiration and ATP generation (sucC, *sucD*, *nuoL*, *nuoG*, *atpA–H*) are likewise suppressed, consistent with oxygen-limited interiors in which oxidative metabolism is constrained and metabolic activity stratifies with depth [73]. Carbon uptake and catabolism loci characteristic of rapidly growing planktonic cells (*glpK/glpX, glcB/glcD/glcG, fucK, dgoT, frlD*) are relatively higher in low-biofilm replicates, aligning with the shift from growth to maintenance observed during biofilm development [11]. Stress-linked and redox-connected enzymes (*putA; gcvP/H/T*) track with this pattern, while *tnaC*, which modulates indole production, shows the inverse association: higher *tnaC*/indole activity corresponds to lower biofilm formation, in agreement with indole’s established anti-biofilm signaling in *E. coli* [74].

Overall, the wild type strain background showed clear replicate-to-replicate divergence, suggesting a moderate degree of stochastic engagement of biofilm programs. Clustering decomposed this variation into antagonistic modules: (i) an adhesion/matrix–on, motility–off program (Cluster 6) centered on EcpR/MatA and reinforced by *nfrB*, *ydeO*, *hipA*, *yhcN/bhsA*, and *nhaA/nhaR*; (ii) a maturation/efflux module (Cluster 1: *sslE*, *ecpC*, *ariR/ymgB*, *mar/MDT*); (iii) a motility/early-attachment/T2SS arm (Cluster 2: *flhA*, *fliD*, *gspM*); and (iv) an inversely correlated growth–respiration program (Cluster 14). Together, these patterns indicate that high-biofilm wt replicas arise when a MatA-centered adhesion circuit, coupled to c-di-GMP–linked EPS synthesis and stress/pH-homeostasis, activates stochastically, whereas low-biofilm replicas retain motility and growth-respiration signatures.

### In Δ*uxuR*, Biofilm Phenotypes Are Strongly Coupled to Repression of Motility and Activation of Acid-Stress Responses

With *uxuR* completely deleted (removing both the UxuR regulator and the *uxuR*-derived sRNAs), this background exhibited the strongest replicate-to-replicate stochasticity among strains. Analysis of six replicas yielded 28 clusters. (Fig. 3).

**Fig. 3.**
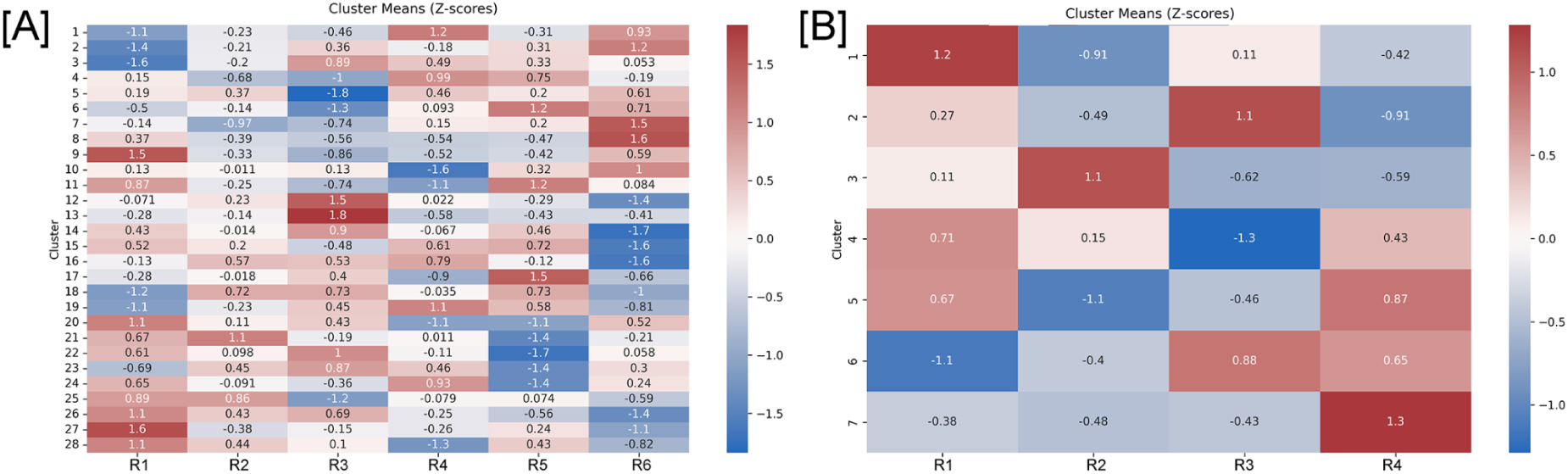
Heatmap of gene expression pattern clusters for *E. coli* K-12 MG1655 *ΔuxuR* [A] *ΔuxuR_*t [B]. Values indicate mean Z-scores, the respective color scale is shown on the right.

In R2/R6 biofilm levels were highest, while in R3 they were comparatively low. The loss of *uxuR* and its intragenic sRNAs, normally capable of dampening motility/chemotaxis (e.g., direct effects on *fliA*), provides a plausible basis for heightened variance in the motile-adhesive switch in this strain [35].

Cluster 13 captured a coherent motility/chemotaxis module that was up in low-biofilm replicas and down in high-biofilm replicas, comprising core flagellar assembly, export, and motor genes (*flgA*–*L*, *flgM*/*N*, *flhB*, *fliA*, *fliC/D/E/F/G/H/I/K/L/M*, *motA/B*) together with chemotaxis components (*cheA/W/R*, *tar/tap/tsr*). Classic genetics established that motility, though not strictly chemotaxis, is required for normal early biofilm formation in *E. coli* K-12, whereas mature biofilms typically repress the flagellar program; our directionality (motility high in low biofilm; reduced in high biofilm) matches this paradigm [71].

Consistent with a motile-state bias in low-biofilm replicas, the c-di-GMP gate was shifted toward low c-di-GMP: the phosphodiesterase *yhjH* (PdeH), which lowers c-di-GMP and thereby sustains motility via the YcgR motor brake pathway, tracked with the low-biofilm state, while diguanylate cyclases (DGCs) that elevate c-di-GMP (and promote curli/matrix) were comparatively inhibited. This aligns with the inverse coordination of motility and curli/biofilm programs in *E. coli* and with the YcgR-mediated motor control [75].

Taken together, Δ*uxuR* resolves into a motility/chemotaxis–dominated Cluster 13 opposed to matrix-promoting, c-di-GMP–high states: in the absence of UxuR and its sRNAs, the network appears more prone to stochastic engagement of the motile program (low c-di-GMP, high flagella) versus commitment to adhesion/matrix, producing the observed replicate dispersion. This interpretation integrates our transcriptional patterns with established flagellar genetics and c-di-GMP logic in *E. coli*.

Cluster 25, on the other hand, is a regulatory/stress module that rises in high-biofilm replicas and falls in low-biofilm ones, consistent with commitment to adhesion/matrix programs and a shift toward slow-growth physiology. The global nucleoid protein H-NS, which modulates surface attachment and flagellar/LPS expression under biofilm-relevant conditions, anchors this module, suggesting broad transcriptional rewiring during surface lifestyle transitions [76]. Small RNAs are prominently represented: OmrA (with OmrB) participates in the sRNA layer that tunes motility and outer-membrane/biofilm programs, including repression of curli regulator CsgD under defined conditions [77]; CsrC antagonizes CsrA to favor curli/PNAG and surface adhesion, and was identified in sRNA overexpression screens as a biofilm-promoting factor [78]. The glucose-phosphate stress sRNA SgrS (and peptide SgrT) also appears here; while its effects are context-dependent, systematic phenotyping showed SgrS overexpression increases biofilm formation in at least some conditions [78]. Matrix processing is represented by *dsbB* (disulfide bond formation), which is connected to cellulose output through a defined, CsgD-independent pathway in UPEC, linking envelope oxidative folding to extracellular matrix composition [79]. Finally, *rmf* (ribosome modulation factor) is a marker of ribosome hibernation and persister-like, stationary-phase physiology often enriched in mature biofilms and tolerant states [14]; and *hdeA*/*hdeB* (periplasmic acid-stress chaperones) plausibly reflect adaptation to pH gradients that develop within biofilms [2].

Cluster 5, which is similar in the pattern, with high expression in high biofilm replicas and low in low biofilm, is enriched for stress-response and envelope-maintenance functions that plausibly reinforce a mature, slow-growth biofilm state. The AraC-family regulator ydeO, an upstream activator of the gadE acid-resistance master regulator, anchors an acid-stress module frequently induced alongside σ^S programs in surface-attached communities; genome-level studies place YdeO within the EvgS/EvgA – YdeO – GadE cascade that drives glutamate-dependent acid resistance [80]. Consistent with oxidative and redox gradients in biofilms, rsxB (of the rsxABCDGE SoxR-reducing complex) marks reinforcement of redox homeostasis that resets oxidized SoxR and tunes the SoxRS regulon after transient oxidative stress [81]. Efflux capacity is represented by emrY (EmrKY-TolC), whose operon is directly activated by EvgA and contributes to multidrug tolerance and envelope stress adaptation, features repeatedly linked to biofilm robustness [82]. Envelope polysaccharide biogenesis is reflected by wecF (enterobacterial common antigen, ECA); although ECA’s biofilm role can be context-dependent, ECA and other surface glycans modulate surface properties and stress resistance, and EPS/LPS pathway perturbations are known to alter E. coli attachment and biofilm architecture [83]. Finally, *yhiD* and *yibS* fall within σ^S-associated stress regulons frequently enriched in K-12 biofilm transcriptomes, consistent with stationary-phase-like physiology in high-biofilm states [84].

Overall, deleting *uxuR* along with its intragenic sRNAs yielded the strongest replicate-to-replicate stochasticity among strains, consistent with loss of a buffering input to the motile-adhesive switch. Clustering resolved this dispersion into antagonistic programs: a motility/chemotaxis axis (Cluster 13) that predominates in low-biofilm replicas and is coupled to a low c-di-GMP state (high yhjH; DGCs subdued), versus high-biofilm stress/regulatory modules (Clusters 25 and 5) comprising global and sRNA regulators (H-NS, OmrA/OmrB, CsrC, SgrS), envelope/matrix maturation (DsbB, ECA/wecF), efflux (EmrKY), and slow-growth acid/redox defenses (YdeO, Rsx, RMF, HdeAB). These patterns indicate that, without UxuR control, the network more readily toggles stochastically between a motile attractor (flagella high, c-di-GMP low) and an adhesive/stress-adapted attractor (matrix and σ^S-linked programs high). The clustering thus turns apparent “noise” into structured modules, identifies likely drivers of state switching (e.g., yhjH vs DGCs; H-NS; CsrC/OmrA; EmrKY/DsbB; YdeO), and provides concrete intervention points to bias Δ*uxuR* cultures toward or away from high-biofilm commitment.

### Translationally Inactive *uxuR* with retained sRNAs Exhibits Reduced Biofilm Variability

Disrupting *uxuR* translation while preserving transcription and locus-derived sRNAs markedly attenuated replicate-to-replicate divergence relative to Δ*uxuR* (Fig.3 B) In the t strain, 4 replicas were studied with genes grouped into 7 clusters. Replica 3 had the slightly higher biofilm mass, with Cluster 2 gene expression elevated in this replica, while Cluster 4 gene expression was comparatively low.

Overall, in respective clusters we observed the co-activation of DgcC-driven c-di-GMP signaling [75] fimbrial adhesion (Sfm), and envelope/redox/stress adaptations (Ivy, Znu, Osm, Dsb/Ccm, fumarate respiration, glycogen, LamB), while the opposing growth/respiration program (TCA, NDH-1/ATP synthase, sugar/glycerol uptake) was concurrently suppressed. Crucially, the overall muted replica dispersion, in sharp contrast to Δ*uxuR*, supports a buffering role for the *uxuR*-derived sRNAs, which appear to constrain bistable transitions between motile (low c-di-GMP) and adhesive (high c-di-GMP) attractors, yielding a more stable transcriptional landscape in this strain.

### Deletion of *crp* Uncovers a Biofilm Module Anchored by PNAG Export and Stress Adaptation

In the Δ*crp* background 4 studied replicas yielded 3 clusters of genes differing in expression patterns.Biofilm level was highest in R1 and lowest in R3 (Fig. 4). Mechanistically, CRP is a global carbon sensor that (i) activates the flagellar master regulator *flhDC*, thereby indirectly promoting motility, and (ii) represses the sRNAs *csrB*/*csrC* (directly for *csrC*, indirectly for *csrB*), which antagonize the RNA-binding protein CsrA. Since CsrA post-transcriptionally represses the *pgaABCD* operon (PNAG biosynthesis), loss of CRP is expected to reduce motility potential and, via the Csr system, de-repress PNAG production, biasing cells toward adhesion/matrix commitment [30,31].

**Fig. 4.**
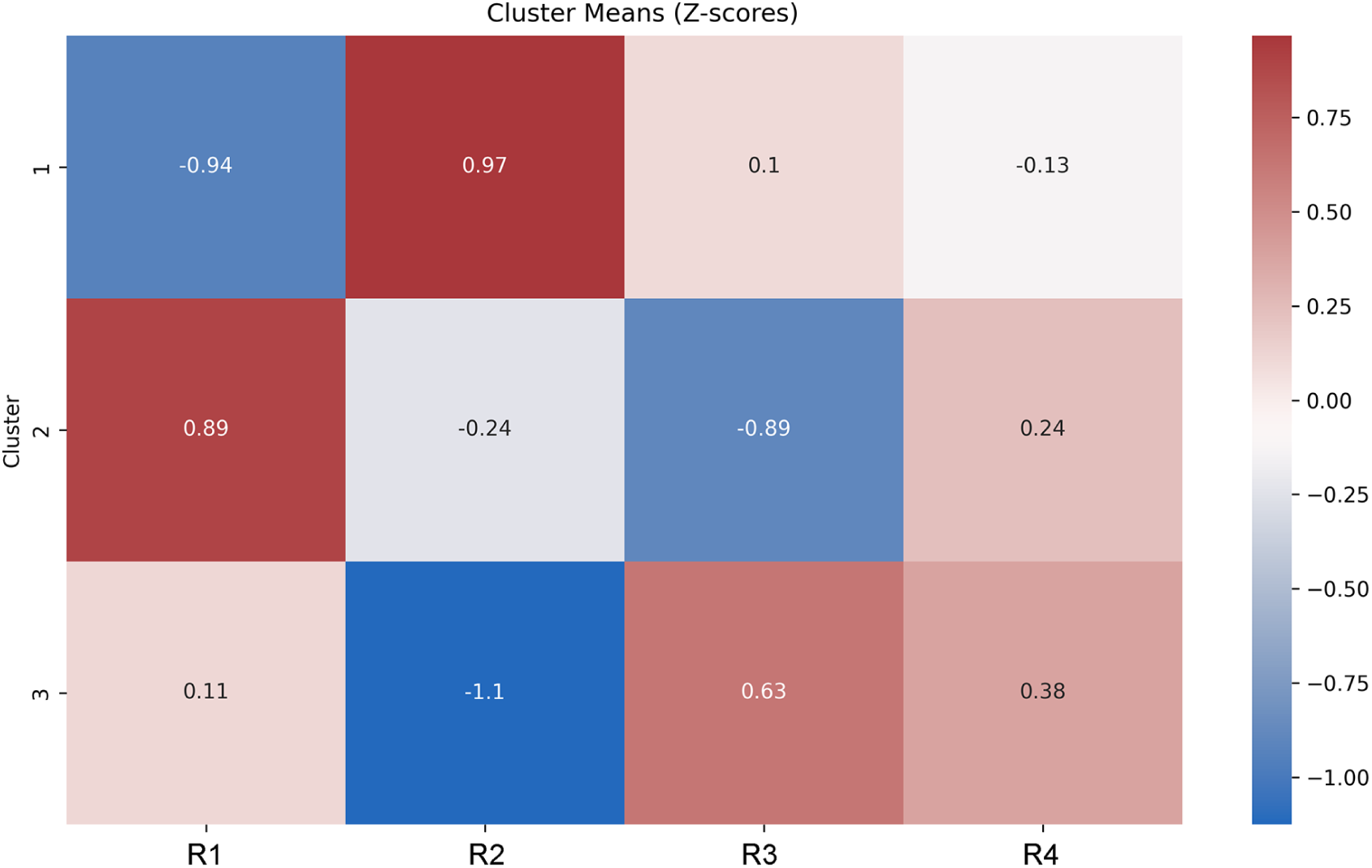
Heatmap of gene expression pattern clusters for *E. coli* K-12 MG1655 *Δcrp* [A]. Values indicate mean Z-scores, the respective color scale is shown on the right.

Genes belonging to Cluster 2 demonstrate high expression level in high biofilm replicas and vice versa and point to a matrix-on, stress-adapted program in the absence of cAMP–CRP control. Cluster 2 contains the direct matrix determinant *pgaA*, which encodes the outer-membrane exporter for PNAG, a polysaccharide adhesin essential for robust *E. coli* biofilm formation and intercellular cohesion [30]. Stress/physiology genes classically enriched in mature biofilms co-occur in this module: *nhaA* (Na⁺/H⁺ antiporter; pH/ion homeostasis), osmC (osmotic/oxidative stress protein), and *cstA* (carbon-starvation protein), consistent with the slow-growth, stress-adapted physiology typical of high-biofilm states [30,62,85]. Metal balance is represented by *rcnA* (Ni/Co efflux), aligning with the metal limitation/redistribution frequently reported inside dense communities [2]. Envelope remodeling potential is suggested by *murR* (regulator of peptidoglycan recycling), which interfaces with cell-wall turnover processes often altered during biofilm maturation, even though *murR*-specific biofilm knowledge in K-12 remains limited. [86].

Deletion of *crp* led to de-repression of *pgaA*, uncovering its clear connection to biofilm levels. Together with overall low biofilm variation level in the *Δcrp* strain and known CRP indirect influence on motility, these results confirm that CRP shapes biofilm phenotypes by “regulating the regulators”.

This work was supported by RSF 24-24-00435.

## Notes

### Competing Interest Statement

The authors have declared no competing interest.

